# A Combined Computational Fluid Dynamics Modeling and Geometric Morphometrics Methods Approach to Quantifying Hemodynamic and Anatomical Features of Embryonic Chick Heart Anatomies Reconstructed from Light Sheet Fluorescence Microscopy Imaging

**DOI:** 10.1101/2024.10.08.611246

**Authors:** Kirsten Giesbrecht, Simone Rossi, Sophie Liu, Shourya Mukherjee, Michael Bressan, Boyce Griffith

**Author notes:** contributed equally (Sophie Liu, Shourya Mukherjee, Michael Bressan).

## Abstract

Although congenital heart defects occur in approximately 1% of newborns in the US annually, their pathogenesis remains largely unknown. Less than a third of congenital heart defects are traced a known genetic or environmental cause. It has been demonstrated that hemodynamic forces such as wall shear stress are critical for heart development. However, measuring these hemodynamic factors *in vivo* is infeasible due to physical limitations, such as the small size and constant motion of the embryonic heart. An alternative approach is to recapitulate the hemodynamic environment by simulating blood flow and calculating the resulting hemodynamic forces through computational fluid dynamics modeling. We use computational fluid dynamics modeling to quantify hemodynamics in a cohort of cell-accurate embryonic chick heart anatomies reconstructed using light sheet fluorescent microscopy. Additionally, we perform a quantitative analysis on geometric features using geometric morphometric methods. Together, the high-resolution but accessible imaging technique of light sheet fluorescence microscopy to reconstruct the anatomies paired with computational fluid dynamics modeling and geometric morphometrics methods produces a fast and accessible pipeline for quantitative hemodynamic and anatomical analysis in embryonic heart development.

## INTRODUCTION

Congenital heart defects (CHDs) are the most common cause of birth defects in humans^1^ and occur in approximately 1% of newborns in the United States^2^. Despite their prevalence, only 20-30% of CHDs can be traced to genetic or environmental causes^3–5^. It is becoming increasingly clear that primary alterations in cardiac biomechanics and hemodynamics significantly impact proper morphogenesis of the heart and developing vasculature. Indeed, phenotypes similar to human CHDs can be robustly reproduced by cardiac outflow tract banding or vasculature ligation in embryonic model organisms^6,7^, suggesting perturbed hemodynamics contribute to abnormal cardiovascular development and function. Particularly, these surgical studies have shown that shear stress regulates several aspects of cardiogenesis through mechanosensitive pathways^8,9^.

Although the relevance of factors such as shear stress is becoming increasingly emphasized as critical for heart development, obtaining accurate shear stress measurements and/or experimentally determining how shear stress regionally varies are both prohibitively difficult due to the minuscule size, delicate nature, and continuous pumping of the embryonic heart^10^. Thus, we currently lack experimental tools that can quantitatively describe spatial and temporal intracardiac hemodynamics which precludes direct modeling of how perturbations to these features would contribute to altered development. Computer modeling approaches for assessing fluid forces, such as computational fluid dynamics (CFD) simulations, offer tractable alternatives to direct experimental measurements. To date, however, a major barrier to the broad application of such CFD models to cardiovascular developmental research has been the lack of protocols for generating cell-accurate descriptions of the embryonic cardiovascular anatomy through which to run simulations. Herein, we use simulations of blood flow and the resulting shear stress throughout the heart and surrounding vasculature to capture spatial and temporal patterns of shear stress^11^ informed by cell-accurate imaging of a cohort of embryos.

Generating cell-accurate three-dimensional geometric data of a cohort of embryos poses new challenges for unbiased geometric analysis compared to traditional microscopy imaging analysis and requires harnessing tools than have not been traditionally used to examine phenotypic differences in developmental biology imaging data. There is no consistent method in developmental biology to analyze volumetric data quantitatively. However, other fields, including paleontology, evolutionary biology, and anthropology, have used morphometrics to evaluate three-dimensional spatial data since the 20th century^12^. Morphometrics is an application of statistics for evaluating specimen shape and size^12^ using data such as lengths and angles of structural features for analysis^13^. By the 1980s, geometric morphometrics emerged from traditional morphometrics methods to quantify more complicated anatomical features, such as vertebrate skulls, using a landmark-based approach^12,13^. Geometric morphometrics has been used in a few developmental biology studies, such as examining phenotypic response to gene expression manipulations ^14^, but it has not yet been widely adopted by the field.

In geometric morphometrics methods (GMM), landmarks or pseudolandmarks defined using Cartesian coordinates are assigned as homologous reference points across a group of specimens and used to compare geometric similarities and differences among specimens in a sample^15^. There are various GMM approaches, with the most common being the Procrustes method or generalized Procrustes analysis (GPA) in biological fields^12,13,16^. The GPA involves standardizing landmark position, scale, and orientation for each set of landmark coordinates for every specimen by translating, scaling, and rotating the landmarks to a common centroid using a least squares approach between each specimen and a mean shape^12,17^. Dimensionality reduction techniques, such as Principal Component Analysis (PCA), are often used in morphometrics analyses to visualize results of highly complex multidimensional datasets produced from the GPA. Visualization methods such as plotting the mean shape and plotting PC scatter plots can be used to interpret the biological results of the data^12^. Here, we apply the GPA to evaluate similarities and differences that emerge across a cohort of embryonic chicks to understand what natural variation is expected and to illustrate GPA sensitivity to geometric differences.

Chick embryos are attractive experimental models for studying normal and pathological blood flow in cardiogenesis^18–24^ due to their similar order of developmental events^20,25^ and size to the human fetal heart, as well as their ease of manipulation and visualization^26,27^. Embryonic chick hearts have previously been modeled in CFD studies^18,28–30^. However, most chick embryonic heart CFD studies of intracardiac hemodynamics employ idealized geometries or combine several individual anatomies to form a single representative geometry of a developing chick heart^31,32^. Comparatively few studies reconstruct individualized anatomies from distinct embryonic chick hearts, but these cohort-based studies have reported that areas that display anatomical variability across embryos do impact local shear stress. Unfortunately, this level of detail is often lost in subject-averaged models^28^, and different methods of constructing local tissue geometries have yielded wide-ranging data on a host of basic morphological features such as average aortic arch midpoint diameter. These studies also report a broad range of shear stress patterns (Table 1).

**Table 1.**
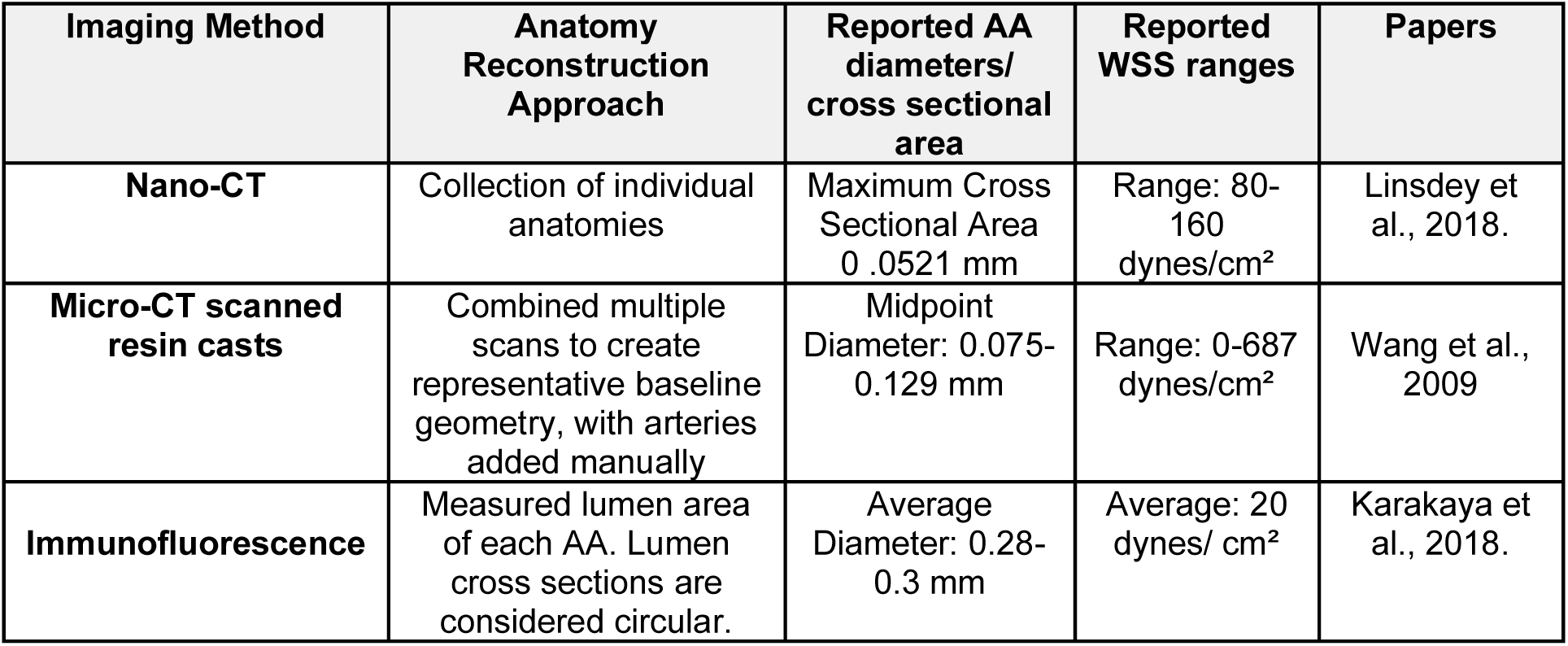
Reported imaging methods and results of hemodynamic and geometric measurements in HH18 embryonic chick AAs.

We chose to simulate blood flow through the aortic arches and the outflow tract, as these are traditionally difficult to image regions, and are prone to congenital heart defects^18,25,33,34^. Most congenital heart defects are concentrated in the outflow tract^18^, a transitory region that connects the early heart to the aortic arches (AAs)^35^ and eventually gives rise to the pulmonary trunk and aorta. Congenital defects occurring in the outflow tract have a high morbidity and mortality rate^35^. The AAs undergo significant remodeling during embryonic growth and contribute to the aorta and pulmonary vein^29^. Generally, 3-4 pairs of AAs are present at a time, and 3 pairs are present upon completed cardiac development^36^. Because they undergo extensive remodeling during development, the arches are prone to developing severe congenital heart defects^34^. The outflow tract and AAs are sensitive to hemodynamic perturbations^25^ and are the sites for half of the congenital heart defects in newborns^25,33^.

Thus the aim of this study was to develop a robust and accessible biocomputational pipeline to capture and compare variations in geometric features and resulting shear stress distributions within intracardiac regions prone to congenital heart defects^35^. Specifically, we designed this pipeline with the intent to construct high-resolution, cell-accurate descriptions of local anatomy in which analysis of flow dynamics could be rapidly performed across cohorts of healthy or perturbed cardiac anatomies (Figure 1). To accomplish this, we have combined techniques for microinjection-based intravital labeling of the embryonic vasculature with light sheet microscopy. We then extracted the vascular geometries of these embryos to simulate blood flow through them. To our knowledge, this is the first study utilizing light sheet fluorescence microscopy to reconstruct subject-specific embryonic chick heart anatomies for computation simulation. This is significant because the relative ease of use and availability of light sheet imaging systems promise to enable high-throughput studies of anatomical variation in large cohorts, largely eliminating the need to perform simulations with idealized geometries. Herein, we use the full incompressible Navier-Stokes equations to simulate blood flow. We numerically solve these equations using the finite element method, which is particularly well-suited for models involving complex geometries^37^. Additionally, we use the GPA morphometrics method to unbiasedly examine geometric variation across the cohort of three-dimensional geometries, including geometries of specimens with less than six AAs. Together, the techniques described below provide a powerful new system to evaluate previously challenging hemodynamic features of the embryonic heart.

**Figure 1.**
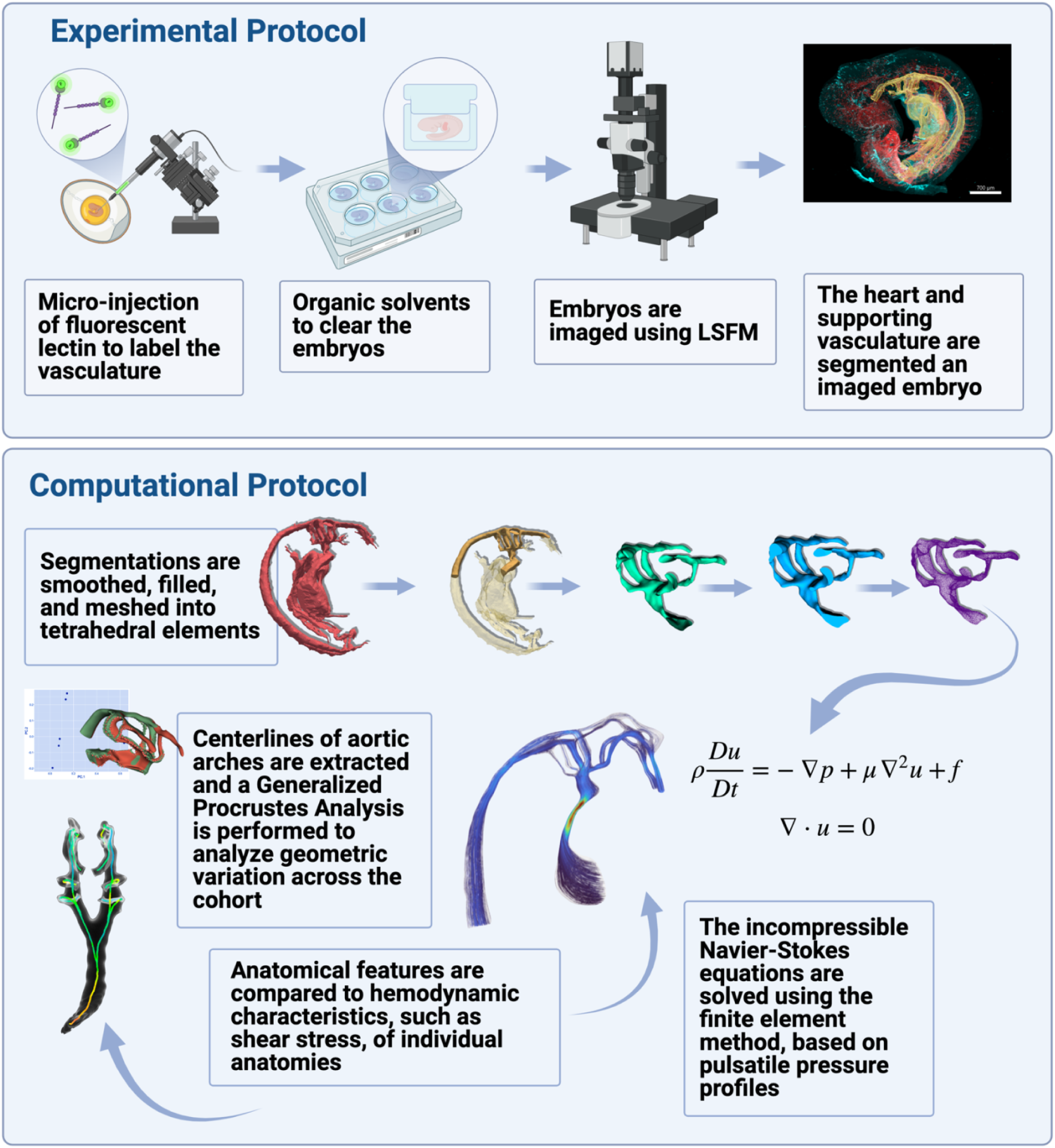
Overview of combined experimental and computational pipeline to compare hemodynamic environments and shape variation across a cohort of embryonic chick hearts. Lectin-tagged fluorophores were injected into the extra embryonic vitelline vein of chick embryos. Embryos were placed in agarose containers and cleared following the iDISCO+ protocol with alternative pretreatment. Cleared embryos housed in agarose boxes were imaged using light sheet fluorescence microscopy. Embryo heart and vessel reconstructions were segmented from the images using Imaris image analysis software. Segmentations were cropped, repaired, solidified and meshed. The GPA was performed across the cohort of embryos to determine how embryos varied geometrically. Centerlines were extracted from the segmentations to calculate geometric features such as radii and length of aortic arches of each embryo. Blood flow described using the incompressible Navier-Stokes equations was simulated through the anatomies by the finite element method. Anatomical variation and hemodynamics were compared across the cohort.

### Reported HH18 Embryonic Chick Morphological and Hemodynamic Properties

## RESULTS

### Intravascular Lectin Injections

To inform the fluid domain of the CFD model we sought to build a ‘cell accurate’ geometry of regions within the heart and the proximal vasculature. In addition, we wanted to utilize a technique for generating this geometry that was adaptable to various experimental techniques (such as outflow banding or vascular ligation) and was compatible with live-imaging approaches. Due to these criteria, we selected live cell labeling using *Lens culinaris* agglutinin (LCA). LCA is a member of the lectin family of carbohydrate-binding factors that labels glycosylated moieties presented on the surface of endothelial cells. LCA has a high affinity for chick vascular endothelial cells at early stages of development, when sensitivity to blood flow perturbations on cardiac morphogenesis is believed to be high^38,39^, and can also be used at later stages of chick development^40^. This means our labeling protocol can easily be adapted to any stage of development. Further, unlike other lectins, including wheat germ agglutinin (WGA), LCA has the ability to bind to finer vessels, appropriate for this early embryonic stage and resolution we aimed to capture in imaging^40^. To increase utility for deep tissue imaging, we utilized LCA conjugated to the far-red fluorescent fluorophore DyLight 649. For the applications noted in this report, we focused on generating vascular geometries from Hamilton Hamburger (HH) stage 16-17 embryos (approximately embryonic day 2.5). Herein, LCA (1:10 diluted in Hanks Buffered Salt Solution) was back-loaded into a pulled glass microcapillary Glass capillaries (1.0 OD/0.7, World Precision Instruments) and 2-5 µl were pressure injected into circulation via the extraembryonic vitelline vein. A protocol of pulsed injections (200 kPa) using a FemtoJet Microinjector (Ependorf) was used to deliver LCA. Using this approach, vascular and heart labeling can be immediately confirmed using a standard fluorescent dissecting microscope.

### Clearing Whole Embryos Housed in Agarose Containers for Imaging

Based on the overall tissue dimensions that we sought to digitally reconstruct for CFD simulations, we selected light sheet fluorescence microscopy as a capture method due to its speed and high resolution^41,42^. Light sheet fluorescence microscopy also minimizes photobleaching and light scattering compared to alternative imaging techniques such as confocal microscopy^41^, and is better at imaging soft tissue when compared to micro-CT^43^. While light sheet microscopy has been proposed to be used in CFD studies in zebrafish hearts^44^, currently it is not routinely implemented for this purpose.

To prepare samples, we utilized a modified clearing technique to preserve LCA staining. Unlike most traditional microscopy methods which require placing the sample on a slide or coverslip to image, light sheet fluorescence microscopy requires placing the sample in a fluid-filled chamber^45^. To stay anchored in the chamber, the sample is first secured in a cradle using a screw. The cradle is then placed in the chamber. If samples are too small to be anchored using the screw, such as the embryos in the study, samples are alternatively embedded in agarose^46^. However, agarose embedding has the potential to distort cardiac and vascular geometry. Therefore, we constructed hollow agarose boxes with removable lids to house each embryo during clearing and imaging. Each embryo was placed in an agarose box, secured with agarose lids molded from custom 3D-printed molds designed in Fusion360 (Autodesk)^47^. After placing the embryos in the lidded, agarose containers, we used the iDISCO+ organic solvent-based clearing protocol with the Alternative Pretreatment^48^ to clear whole embryos. We selected this iDISCO+ protocol as the clearing method due to its low cost, speed for clearing embryo tissues, and success at clearing tissue imaged using light sheet fluorescence microscopy, compared to other clearing methods^49^.

### Imaging Whole Embryos Using Light Sheet Fluorescence Microscopy

After being cleared, embryos were imaged on a Lavision BioTec Ultramicroscope II light-sheet system. The pixel size was between 1.2-1.95 µm, with each Z slice spaced 5 µm apart and a horizontal focus of 5 µm. To excite the stained lumen and autofluorescence, a 647 nm laser excitation and 488 nm laser excitation were used to image two channels, respectively, for each embryo. The Imaris (Bitplane, Oxford Instruments)^48^ File Converter software was used to convert light sheet fluorescence microscopy-generated TIFF files to Imaris files.

### Image Segmentation

Following image acquisition, cardiovascular anatomy must be converted to a format usable for CFD modeling. For these purposes, we used Imaris software (Bitplane, Oxford Instruments)^50^ to visualize and manually segment the embryonic heart and vasculature geometries. The LCA 649 fluorescence channel combined with the autofluorescence channel were used to define vascular and cardiac lumen structure (Figure 2A-C). As shown in Figure 2 (Figure 2D-F), this resolves fine details related to vascular bifurcations, microvessel anatomy, and cardiac chamber arrangement. As the cardiac outflow tract and AAs have previously been examined using CFD models, we focused on these areas of the heart to confirm the viability of our approach. Typically, imaging the AAs is difficult due to their complicated geometry, small size, and dorsal/anterior position relative to the looping heart, so alternative methods such as ink injections have been used to map the arches^32^. However, we observed high fidelity and reproducibility across samples when imaging these regions with light sheet fluorescence microscopy. Additionally, LCA 649 labeling allowed us to generate high-resolution segmentations of both the outflow tract and AA despite their complex configurations and to clearly distinguish the lumen of the outflow tract and developing chambers from soft tissue within the heart (Figure 2D, E).

**Figure 2.**
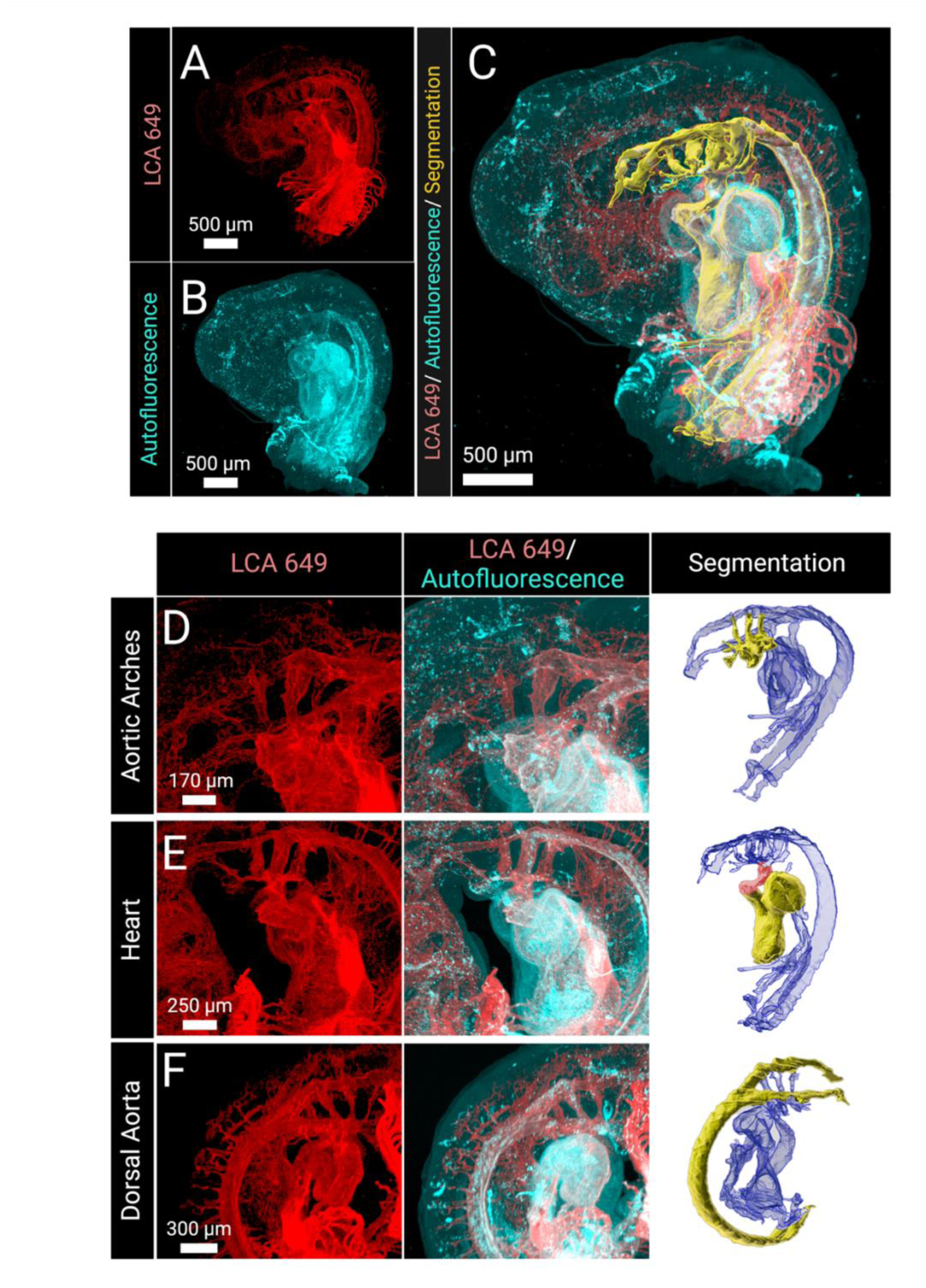
Light sheet fluorescence microscopy optically sections whole embryos for generating anatomy reconstructions. (A) The LCA 649 channel distinguished endothelial cells lining the lumen and was primarily used to inform segmentations. (B) The autofluorescence channel distinguished soft tissue and was used to verify segmentations. (C) Together these channels were used to construct a segmentation of the developing heart, AAs, and dorsal aorta. (D-F) The imaging captured anatomical features that are often difficult to image, due to the size and position of these features. In the segmentation panel, the segmented portions of interest are highlighted as solid yellow segmentations within the entire segmented anatomy, which is shown in blue. (D) AAs, which are notoriously difficult to image due to their small size and position were segmented using the LCA 649 channel. (E) The heart, including the outflow tract was segmented. The LCA channel distinguished the lumen from other soft tissue, such as the cardiac jelly, in the heart which can be seen in the autofluorescence channel. The outflow tract is highlighted red in the segmentation. (F) The bifurcating dorsal aorta was segmented using the LCA 649 channel as well.

To translate raw imaging data into three-dimensional geometries appropriate for CFD modeling, we generated a processing pipeline to build structural scaffolds that matched the vascular anatomies obtained by light sheet imaging (Figure 3A). As an example of this process, we extracted aortic arch geometric patterns described above and prepared a cohort of these samples for CFD simulations. Initially, we cropped to a region of interest containing the aortic arches, outflow tract, and dorsal aorta. Then, each “hollow” vasculature structure was post-processed in the software MeshMixer (Autodesk)^51^ to resolve rough boundaries remaining from manual segmentation. Next, the hollow segmentations were translated into solid structures in Blender (The Blender Foundation)^52^ using the Quad Remesher plugin (Exoside)^53^ which retopologizes the geometry to make it suitable for meshing. Following this step, the AAs, outflow tract, and dorsal aorta were meshed and refined using Cubit (Coreform)^54^ into at least 100,000 second-order tetrahedral elements for each anatomy to be used for CFD simulations. Grid spacing was approximately 0.009 mm. Upon completion of segmentation discretization, we analyzed the geometric features of each embryo in the cohort to evaluate our geometries in relation to previous reports.

**Figure 3.**
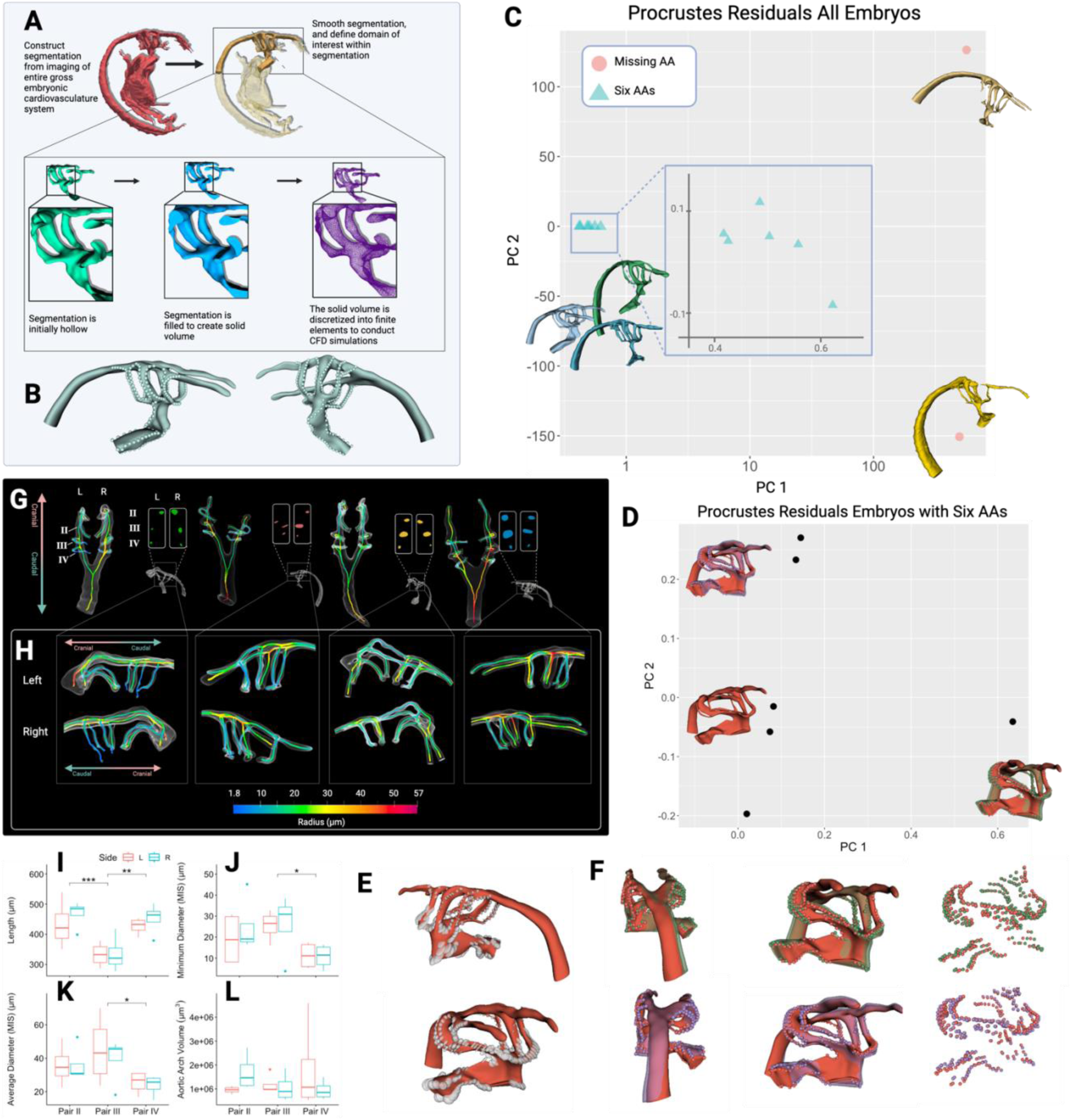
(A) Following manual segmentation from imaging, the region of interest was selected which included the outflow tract, AAs, and dorsal aorta. Next the segmentation was smoothed, solidified and discretized into a tetrahedral mesh. (B) All GMMs are based in landmark or pseudolandmark comparisons. In this study a GPA was performed to analyze shape variation in the cohort. (B) There were 275 pseudolandmarks placed along the contours of geometric features of a template embryo, such as the AAs for the analysis. (C) First, two embryos that were missing multiple AAs were included in the GPA. The PCA scatter plot of the Procrustes residuals separates these two embryos from the rest of the embryos in the cohort, with an inlay plot showing the detail. Note that the x-axis is shown in log scale. (D) Next the GPA was repeated on only embryos containing all six AAs. The PCA plot displays how the embryos varied in PC 1 and PC 2, with the mean shape warped by PC 1 (shown in green) and PC 2 (shown in purple is displayed with the mean shape (shown in red). (E) The variance of each pseudolandmark is plotted as ellipses. The radius length corresponds to the variance in that dimension of an ellipse and shown in two views. (F) The mean shape is warped by PC 1 (shown in green) and PC 2 (shown in purple). The warped pseudolandmarks are also plotted for the respective PCs. Some specimens from the cohort have been highlighted to demonstrate that (G) Aortic arch cross sections are elliptical in shape at this stage. (H) AA III contains the largest average diameter, while AA IV contains the smallest minimum and average diameter. (I-L) Arches were compared laterally (left/right sides) and pairwise (AA II/AA III/ AA IV) using ANOVA followed by Tukey analysis (where * indicates P < 0.05, ** indicates P <0.01, *** indicates P <0.001). In general, arches varied by pair, but not laterally (left vs. right). Diameters were calculated by measuring the diameter of the maximally inscribed sphere (MIS) at a given point in the arch. (I) The central arch, AA III was significantly shorter than either of the other arches flanking it. (J) AA IV had the smallest minimum diameter

### Geometric Morphometrics Analysis

We performed a GPA on the cohort of embryos which included two embryos that lacked at least one AA or cranial arch due to tissue tearing during processing and imaging to examine if the GPA could distinguish embryos with missing AAs from other embryos in the cohort. Using the software Slicer^55^ and the Slicer extension SlicerMorph, We placed 275 pseudolandmarks defined using curves along geometric features such as the AAs and contours of the outflow tract along a randomly chosen template specimen from embryos containing six arches (Figure 3B). After the pseudolandmark set had been defined on the template specimen chosen from a control case, the set was transferred from the template model to all the specimens using the SlicerMorph module ALPACA (Automated Landmarking through Point Cloud Alignment and Correspondence Analysis).

After generating pseudolandmarks for all specimens, we performed the GPA and preserved Boas coordinates to maintain size variation using the corresponding module in the GPA module in SlicerMorph. We observed that the embryos containing all six arches clustered together and were separated from the embryos lacking at least one arch in the PCA scatter plot of the Procrustes residuals (Figure 3C), demonstrating the GPA is well-suited to distinguish gross anatomical differences among specimens.

Next, we removed the outliers containing the missing AAs and repeated the GPA with the six remaining embryos and plotted the PCA scatter plot of the Procrustes residuals. We visualized these results by plotting the mean shape resulting from the GPA and warped the mean shape by PC 1 and PC 2, separately. The mean shape and warped mean shape are displayed on the PCA plot, indicating how the specimens vary with respect to PC1 and PC 2 (Figure 3D). Figure 3E shows the relative variance of each pseudolandmark in each dimension as indicated by the radius length of the corresponding dimension of ellipses corresponding to each pseudolandmark. Figure 3E indicates that relative variance was not uniform across the pseudolandmarks, with pseudolandmarks varying greatly along the outflow tract and left AA II. In Figure 3F the mean shape, indicated in red, warped by PC 1, shown in green, indicates that PC 1 corresponds to a shift in the outflow tract to the left and a twisting about the AAs which resulted in the dorsal aorta shifting to the right. Additionally, PC 2, shown in purple, indicates a downward shift and shortening of the outflow tract, resulting in the dorsal aorta rotating to the right. One specimen was more positively associated with PC 1 compared to the other specimens considered. This suggests that GPA can provide an unbiased method of distinguishing outliers in three-dimensional analysis for similar cohort-based studies. The biological interpretation of the PCA plot of the Procrustes residuals indicates that specimens are likely to vary by outflow tract shape, AA II, and the angle of the dorsal aorta to the AAs. The GPA also demonstrates that shape variance is not uniform across all geometric features in the cohort.

### Basic Geometric and Centerline Analysis

In addition to the GPA, we also analyzed basic geometric features of embryos. We observed that the AA cross-sections are elliptical, generally with the longer axis oriented laterally (left, right) (Figure 3G) and that the central AA III pair typically had the largest diameter (Figure 3H) which is consistent with reports from previous studies that analyze individual embryonic heart anatomies across a cohort from nano-CT imaging data^28^. To measure quantitative geometric features, we extracted centerlines through the dorsal aorta and aortic arches of each embryo (Figure 3G-H) using the vascular modeling toolkit (VMTK) extension^56^ in the open-source image computing platform, 3D Slicer (https://www.slicer.org/)^55^. Comparing AA geometric features by pair and side revealed that AAs primarily varied geometrically by pair, but not by side (Figure 3I-L). The centrally located arch pair, AA III, was significantly shorter than either of its neighboring arches (Figure 3I). The AA IV exhibited the highest degree of stenosis, as measured by the minimum reported diameter across each arch (Figure 3J). The average diameter of AA III was significantly larger than the average diameter of AA IV (Figure 3K). However, the volume of each aortic arch did not vary significantly by arch pairs or sides at this stage (Figure 3L). This pattern that the AA IV arch pair has the smallest average diameter and that AA III has the largest diameter is consistent with previous reports among the diameters of these three arch pairs measured at similar stages^32^.

### Embryonic Heart Anatomical Variability

Following the geometric analysis of each embryonic heart anatomy, we noticed variability in vessel configurations across the embryos. All embryos examined had bilaterally pair aortic arches, but the degree of stenosis in the outflow tracts and AAs varied across embryos. Additionally, we observed a geometry (hereafter referred to as case 1) where the AA II pair had a larger diameter than the AA III. We also observed a geometry (case 2) where there was an outlet in each cranial artery. Section ‘CD simulations’ describes how blood flow varied in these special cases.

### CFD Modeling Parameters

To implement a flexible CFD modeling pipeline that can be used to robustly simulate anatomy specific hemodynamic characteristics, we defined a basic set of blood flow characteristics and pressure profiles for use in cardiac simulation. Initially, we focused on simulating blood flow about the AA anatomies as these vessels showed a high degree of variability across our sample cohort and are particularly susceptible to congenital malformation. Herein, we modeled blood as an incompressible Newtonian fluid with a viscosity, μ = 3.71·10^-3^ Pa·s and density, ρ =1060 kg·mm^-3^ as reported by a previous embryonic blood rheology study^57^. We used the full incompressible Navier-Stokes equations, including the nonlinear convective term. We use the momentum equation to describe blood flow (1) and the incompressibility constraint (2), in which *p* and ***u*** are the fluid pressure and velocity fields, respectively,

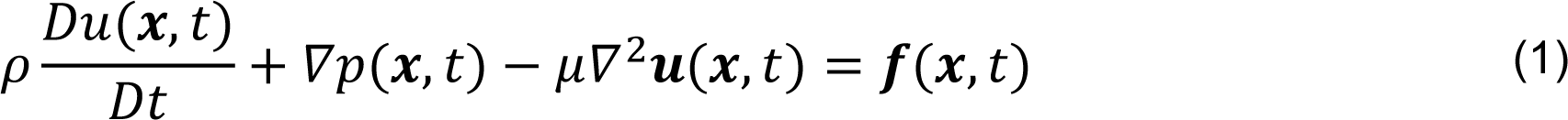

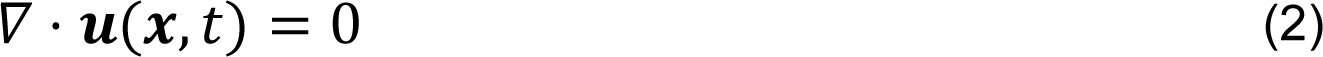

We solved the system of equations using the finite element method because it facilitates models involving complex geometries^37^. The 3D CFD simulations to model blood flow using the finite element method were performed using an in-house C++ code based on the libMesh^58^ finite element library and using linear solvers implemented in PETSc^59^ (see also Materials and Methods). This technique required a mechanism to stabilize pressure^60^. To this end, we used LBB-stable P2-P1 Taylor-Hood velocity-pressure pairs^60,61^. For our simulations, we set the grid spacing to be approximately 0.009 mm, with each anatomy containing at least 100,000 second-order tetrahedral elements. Blood flow was assumed to be laminar^62^. No slip boundary conditions were imposed on intracardiac and vessel walls, which are assumed to be rigid and impermeable. Sinusoidal pulsatile pressure profiles, which we determined by digitizing previously reported pressure profiles in the ventricle and dorsal aorta^63^, were imposed at the inflow and outflow boundaries, respectively. Simulations and resulting hemodynamics were visualized in ParaView (Kitware)^64^.

### CFD simulations

Using the above parameters, we determined how geometric features that varied across our sample cohort impacted hemodynamic patterns within the AAs. We examined WSS both at peak flow (Figure 4A) and during the accelerating phase of blood movement (Figure 4B). As expected, peak WSS occurs during peak flow (Figure 4A-B) and the pressure difference between the dorsal aorta and outflow tract is highest during peak flow (Figure 4F). In general, elevated WSS was observed in stenosed regions of the anatomies. Consistent with our rationale for using a cohort-based approach, we noted that the regions of maximal WSS varied with local AA and outflow tract geometry

**Figure 4.**
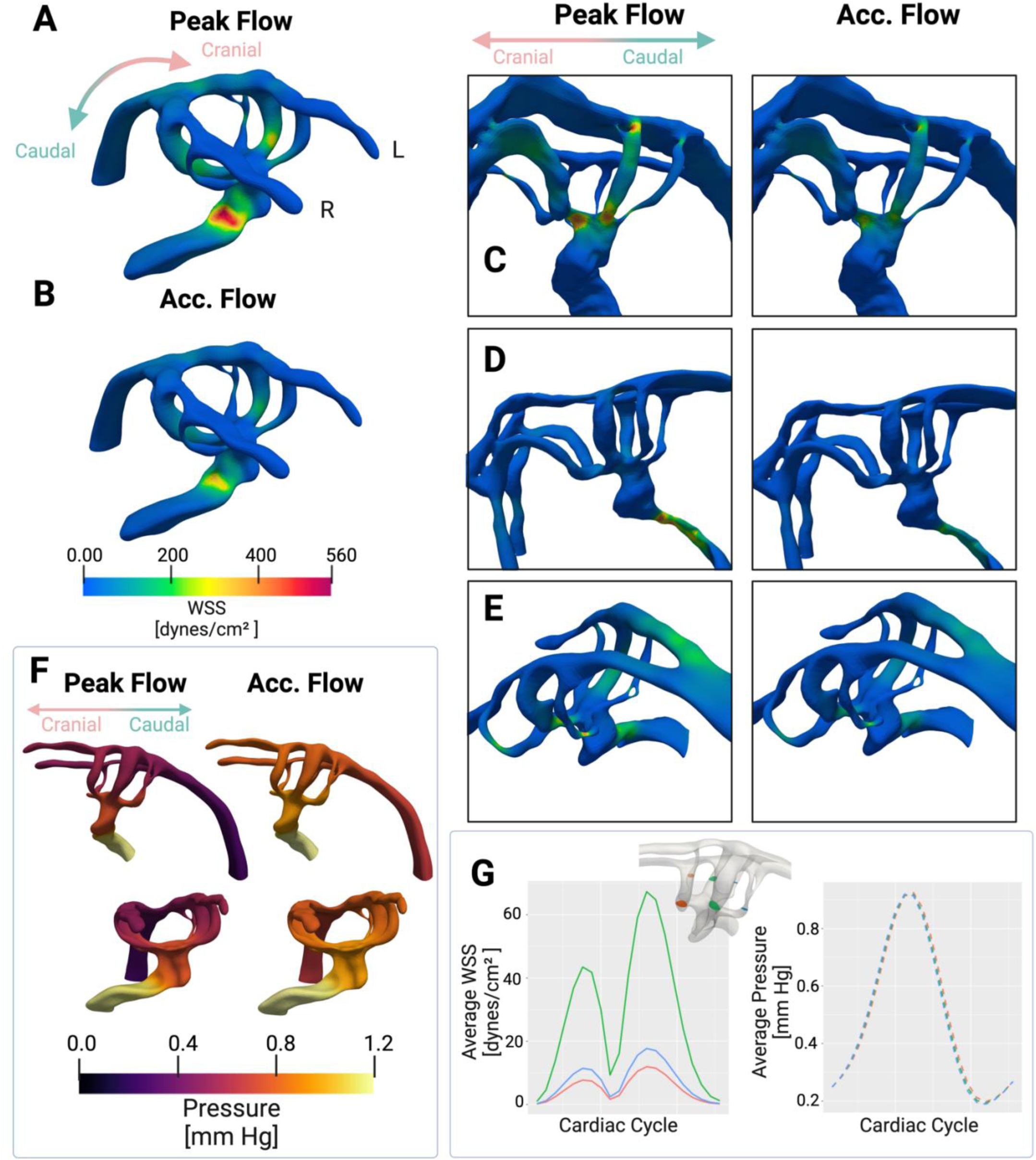
(A) WSS was observed during peak flow and (B) accelerating flow. As expected, peak WSS occurred during peak flow. Peak WSS locations varied by individual anatomical features within embryos. (C) In an embryo (case 1 embryo) with less stenosis in the outflow tract, peak WSS occurred in the AAs. (D) Conversely, in an embryo (case 2 embryo) with a highly stenosed outflow tract, peak WSS occurred in the outflow tract. (E) An embryo with a moderately stenosed outflow tract experienced elevated WSS throughout locally stenosed regions including the outflow tract and AAs. WSS spatial patterning is similar during peak flow compared to accelerating flow. Regions that experience the highest WSS during peak flow experience high WSS during accelerating flow, indicating that this flow regime is not heavily driven by convective effects. (F) The pressure difference between the outflow tract and dorsal aorta was larger between the outflow tract and dorsal aorta during peak flow compared to accelerating flow, as expected. (G) Average WSS was highest in the AA III pair, which is the centrally located pair of the three pairs present during this stage of development, despite all pairs of the AAs experiencing consistent pressure and the higher degree of stenosis in flanking arches of the AA III pair.

(Figure 4C-E). Conversely, if the outflow tract was heavily stenosed, elevated WSS occurred in the outflow tract (Figure 4D). If the outflow tract was moderately stenosed, elevated WSS was observed throughout locally stenosed regions, including the outflow tract and the AAs (Figure 4E). In addition, across embryos spatial WSS patterning was similar during accelerating flow and at peak flow time points (Figure 4A-E). These similar WSS patterns indicate the flow is not heavily driven by convective effects and that the pulsatile hemodynamics of the cardiac cycle do not shift the positions of highest WSS.

An advantage of our CFD modeling approach, compared to an experimental one, is the ability to sample hemodynamic features at any position within the acquired geometry. This allowed us to determine that, surprisingly, the central pair of AAs (AA III) displayed the highest average WSS across our samples (Figure 4G). AA III was the shortest pair present in our embryos, but was not the most stenosed arch pair (Figure 4D-F). Of note, each AA pair experienced similar pressure during the cardiac cycle, despite the central pair experiencing higher WSS (Figure 4F, G). These data indicate stenosis alone of the arches is not a sufficient predictor of WSS within the arches.

A potential explanation for the higher WSS in AA III compared to the flanking arches would be that flow distribution in the arches is not homogeneous. To further examine this, we extracted blood velocities as flow streamlines calculated through the AA anatomies and found that blood flow velocity streamlines were consistent with WSS patterns. Specifically, blood flow velocity was elevated in locally stenosed regions of the embryonic heart where peak WSS was found, such as the outflow tract and AAs (Figure 5A). Elevated velocity was observed in AA III, consistent with the elevated WSS patterns (Figure 4A, 5A). We observed that blood flow velocity patterns, including the locations where peak velocities occurred, were consistent with peak WSS patterns across embryos (Figure 5B-D). The spatial patterning of velocity streamlines also was maintained across acceleration and peak flow (Figure 5). As expected, this is consistent with the spatial WSS patterns maintained during the two different stages of flow. Since the location of peak velocity streamlines is maintained across different flow phases, this further supports the finding that there is little convective mixing.

**Figure 5.**
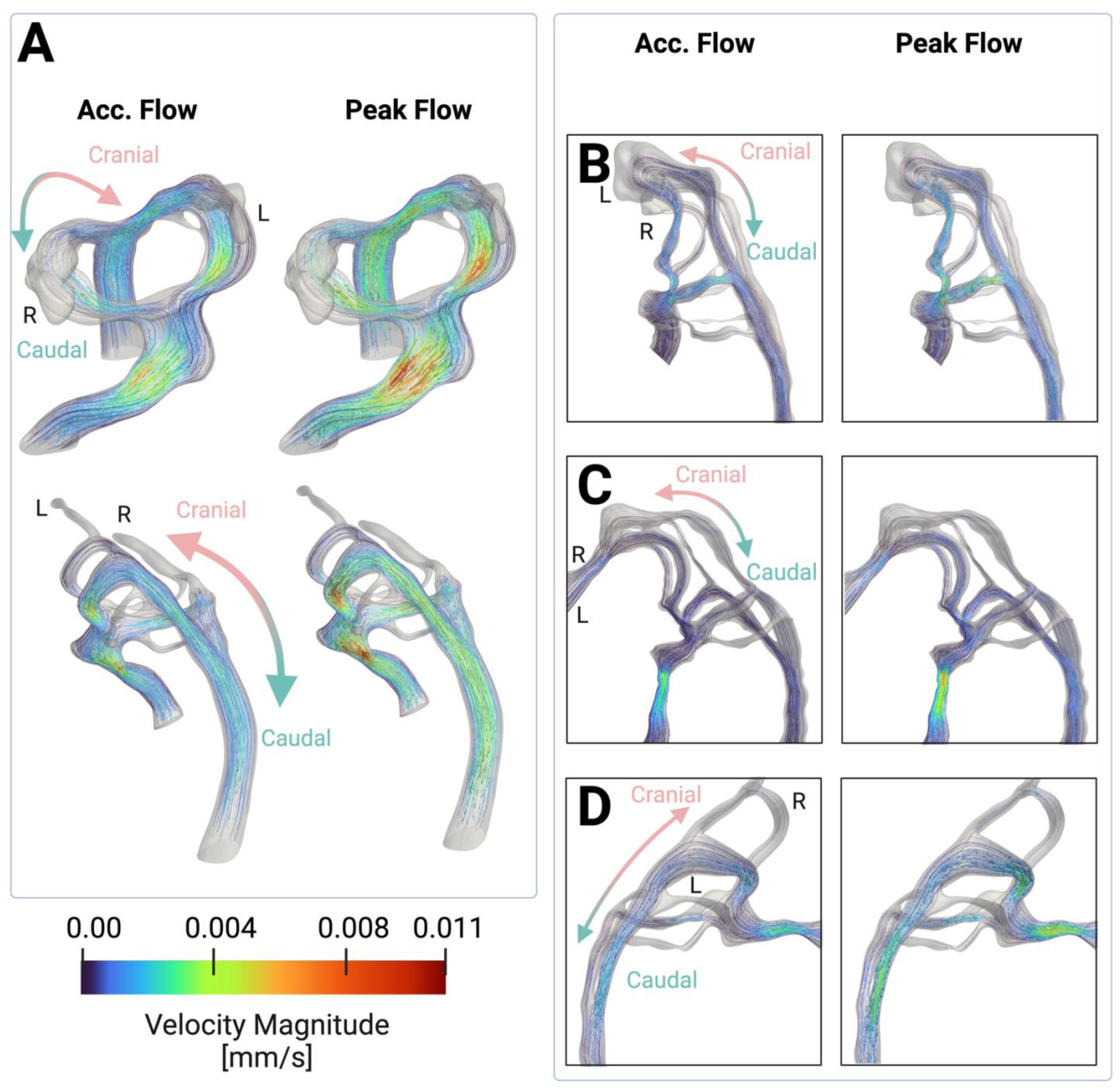
(A) Blood flow velocity streamlines are observed during peak flow and accelerating flow, and as expected, exhibit the similar temporal and spatial patterns compared to WSS, consistent with the finding that there is low convective flow. As expected, blood flow velocity streamlines indicate elevated blood flow velocity in the same regions as elevated WSS across variations in embryonic anatomies. (B) An embryo with little stenosis in the outflow tract experiences elevated velocity primarily in the AAs, consistent with WSS patterns. (C) An embryo with highly stenosed regions in the outflow tract experiences elevated flow velocity in the outflow tract, consistent with the elevated WSS. (D) An embryo with moderate outflow tract stenosis experiences elevated WSS throughout locally stenosed regions in the outflow tract and in the AAs.

To examine how blood traveled through the AAs, we traced streamlines through each AA. We placed a point cloud enclosing each AA cross section to seed each streamline. Next, we traced the streamlines of each AA from the outflow tract to the dorsal aorta. We compared streamline paths from the outflow tract through the dorsal aorta (Figure 6A-D). We observed that the streamlines that flowed through the AA IV pair initially flanked each other in the outflow tract but separated to opposite sides in the cross section of the dorsal aorta after traveling through the arches (Figure 6A-D). Streamlines that contributed to the AA II pair also flanked each other in the outflow tract but did not separate in the dorsal aorta (Figure 6 A-D); instead, they traveled towards the center of the dorsal aorta cross section. Examining the cross section in the outflow tract (Figure 6E) and the cross section of the dorsal aorta (Figure 6F) confirms these patterns. The cross sections of the streamlines demonstrate that flanking AA streamlines in the outflow tract generally maintain their flanked configuration in the dorsal aorta, with the exception of the AA IV paired streamlines which separate. For instance, the R AA II streamlines are flanked by the L AA II and R AA III streamlines in the outflow tract (Figure 6E) and reunite with these two sets of streamlines in the dorsal aorta (Figure 6F). This maintenance of neighbors of streamline sets from the outflow tract to the dorsal aorta was consistent across embryos (Figure 6G).

**Figure 6.**
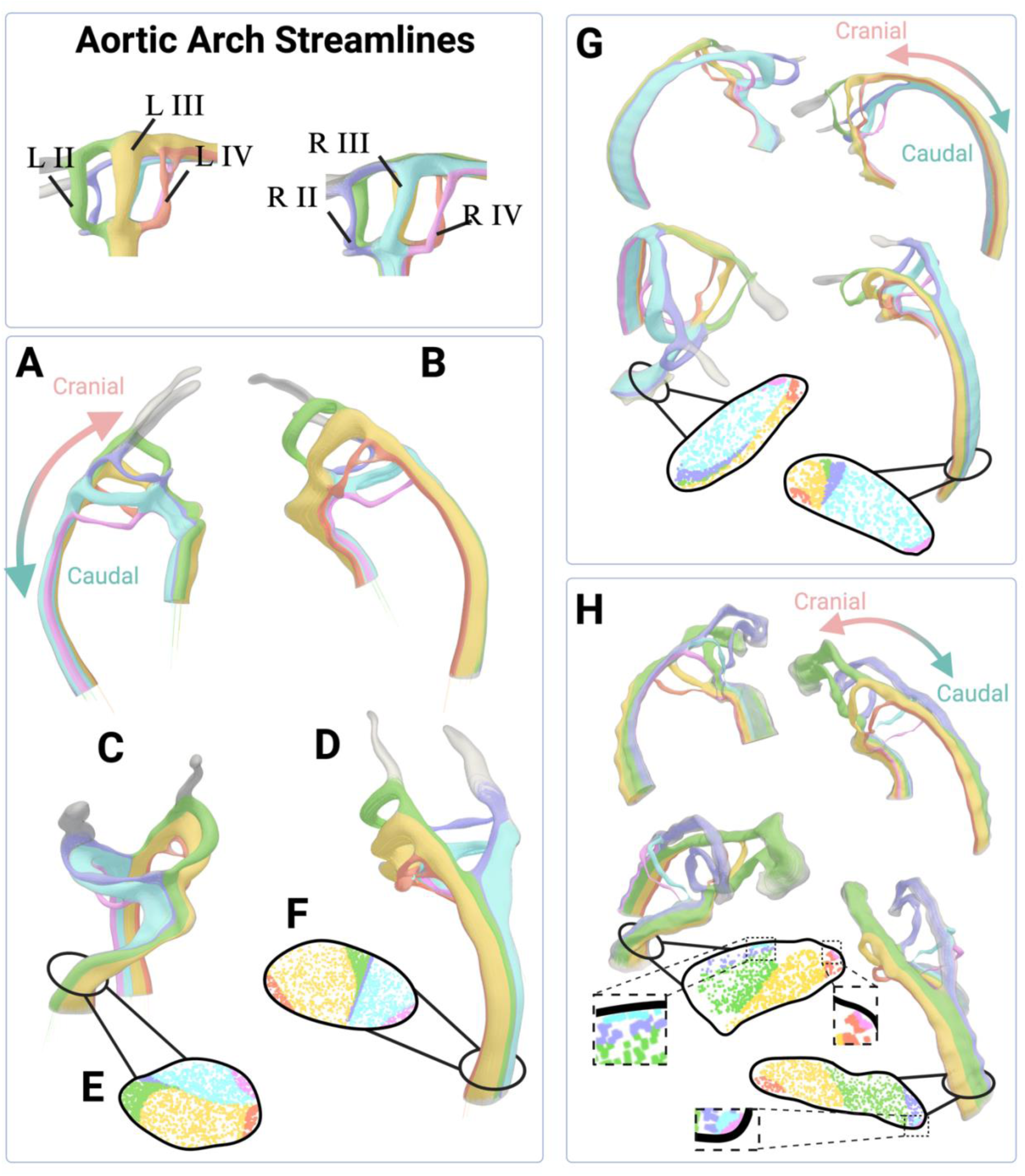
We identified streamlines through each aortic arch. We traced streamline paths from the outflow tract to the dorsal aorta to understand streamline spatial configuration in relation to before and after flowing through the arches. Streamlines through the left AAs (A) and right AAs (B) of an embryo show that there is little mixing of flow streams, even after exiting the AAs. (C) We observe the streamlines caudally and through (D) the dorsal aorta. We observe that the streamlines through AA IV pair are split by the other arch pair streamlines while traveling down the dorsal aorta. (E) We examined streamlines in the cross sections in the outflow tract, upstream of the AAs, (D) and in the cross section of the dorsal aorta, downstream of the AAs. The order of the streamlines in the outflow tract was maintained in the dorsal aorta, and confirms that the AA IV pair separates after flowing though the aortic arches. There is little mixing of streamlines, consistent with the finding that there is little convective influence in this flow regime. (G) A second embryo illustrates that the spatial order of streamlines is largely conserved across anatomical variations. (H) We examine streamlines through an embryo where AA III is smaller than AA II, and show that the distribution, but not order of streamlines is altered and that the order is conserved. Detailed views of the cross sections are shown to highlight all the AA streamlines.

This spatial order of streamlines from the outflow tract to across the dorsal aorta was maintained even in embryos with unusual variation in the AAs. For instance, in case 1, where the AA II pair had a larger diameter than the AA III pair, the dorsal aorta cross section contained the same spatial order of streamlines, despite the difference in flow distribution across the arches (Figure 6H). In case 2, where there were additional outlets in the cranial artery, we observed that the streamlines from the AA II pair contributed to the cranial artery outlets instead of the dorsal aorta outlet. The streamlines from the remaining arches maintained the same spatial ordering in the dorsal aorta cross section (Supplementary Figure 1). These data demonstrate that there is a predictable pattern that streamlines follow before and after entering the aortic arches that is conserved across anatomical variations.

## DISCUSSION

In this report, we have described a fast, accessible, and high-throughput pipeline for reconstructing cell-accurate surfaces of embryonic chick heart lumens that can be used in CFD models of blood flow and geometric analysis. Additionally, we demonstrate that unbiased GMMs can identify embryos that greatly vary phenotypically, as shown in the GPA of the embryo cohort which included embryos that lacked typical anatomical features present at that stage of development. As imaging techniques continue to evolve, highly detailed, multi-dimensional, datasets are becoming the norm. However, standardized techniques for preforming unbiased geometric analysis for conducting morphometric comparisons across sample groups have largely not been identified or implemented. This is particularly problematic in the developing heart, which is a complex, three-dimensional, structure that dynamically changes its size, shape, and alignment across development. Thus, in this report we demonstrate how GMM may address a large gap in the field. Furthermore, we pair the geometric analysis with CFD models to relate specimen-specific geometry to hemodynamic patterns.

We labeled the lumen of the embryos using LCA, which has a higher affinity for embryonic chick endothelial cells throughout development compared to other lectins^40^, such that this pipeline can easily be extended to imaging and assessing embryonic chick geometries and hemodynamics throughout various stages of development. We employed light sheet fluorescence microscopy, requiring only minutes of acquisition time, which provides a fast and accessible imaging method to capture the difficult-to-reach anatomy of embryonic chick hearts. This pipeline could easily be extended to image embryonic hearts in more than two channels, using antibody staining, which is compatible with the iDSICO+ clearing method^48^. Although we primarily focus on region such as the AAs, dorsal aorta, and outflow tract, our imaging results indicate the light sheet is well suited to capture other fine vessels surrounded by soft tissue, and even various soft tissue types. Imaging additional channels could be used to distinguish soft tissues, such as the cardiac jelly and myocardium and could be applied to inform mechanical or structural models of early chick hearts and supporting vasculature. Including additional channels could also highlight flow-sensitive genes to complement CFD studies. Light sheet fluorescence microscopy has already been used to visualize shear stress-activated signaling pathways, such as Notch signaling in embryonic zebrafish hearts^65^. It would be a natural extension and an advantage of using light sheet fluorescence microscopy for imaging to simultaneously examine gene expression and to validate CFD models of subject-specific anatomies.

Our CFD model provides a fast and cheap platform to perturb the hemodynamic environment to predict where changes in blood flow might be greatest in the anatomy. Although we apply this pipeline to a cohort of healthy embryonic chicks, this pipeline could easily be extended to examine how genetic or hemodynamic manipulations can perturb the geometry and the simulated blood flow of embryonic chick hearts at a high visual resolution, particularly in the regions this study identified that experience peak WSS, such as the outflow tract and AAs.

While adult avian and human blood both behave as non-Newtonian fluids, during early stages of development when relatively few red blood cells are circulating, embryonic chick blood is generally considered to be a Newtonian fluid^57^. This study, as well as most CFD studies of these stages and later in chick embryonic development, models blood as a Newtonian fluid^32,66^. However, it may be interesting in the future to examine how modifying the modeled embryonic blood to behave as a non-Newtonian fluid would impact the hemodynamics, particularly the spatial and temporal WSS results.

We numerically solved the full incompressible Navier-Stokes equations using the finite element method, which is well-suited for these complicated geometries. The results demonstrating that WSS is elevated within the outflow tract and AAs may suggest that these regions are especially sensitive to heightened and variable WSS and altered blood flow. These findings are consistent with previous findings that report elevated WSS in the AAs^28^. Additionally, the AAs and outflow tract are prone to congenital heart defects^35^. These findings suggest that perturbed hemodynamics may contribute to congenital heart defects that arise in these regions.

A surprising finding from this study is that radii alone in the AAs was not predictive of peak WSS or blood flow velocity within the AAs. Instead, the pair of arches located centrally, AA III, within the three pairs present experienced the highest WSS. The central pair was also the shortest pair among the pairs present at the stage. These results suggest that geometric features upstream of the arches may influence hemodynamic forces.

From these simulations, we found that there is little convective influence under this flow regime from the consistent velocity and WSS spatial patterns across different time points in the cardiac cycle, and further supported by the little mixing across embryos of arch streamlines in the dorsal aorta after traveling through the arches. To our knowledge, this is also the first report to examine streamlines through each individual arch and trace their distinct behavior downstream and upstream to other arch streamlines.

Additionally, this cohort-based approach revealed elliptical cross sections and aortic arch diameters on the same order of magnitude as AAs imaged in other cohort-based studies^28,67^. Unlike studies that model blood flow through composite or idealized embryonic heart models, which assume circular cross sections, these cohort-based studies are the only studies to report elliptical cross sections for AAs at these stages. In modeling hemodynamics in this system, the elliptical shape could be an important feature to consider for future modeling studies that is often ignored.

The combination of using a cohort-based approach and generating three-dimensional geometric reconstruction allows unbiased and comprehensive three-dimensional geometric analysis to be performed. We use the GPA to unbiasedly identify specimens that lack anatomical features such as AAs. Our findings through the visualization techniques of the GPA results indicate that different geometric features such as the outflow tract and AA II are more likely to vary compared to other geometric features. The GPA also provides an unbiased method to identify outliers for further downstream analysis.

It’s well established that hemodynamics plays a critical role in shaping the course of a developing heart. Unfortunately, technical limitations that prevent direct measurements of blood flow patterns within animal model hearts have stagnated the progress to elucidating the role of hemodynamics in cardiovascular development. This flexible and customizable pipeline overcomes this challenge by using high-resolution imaging to inform CFD modeling simulating blood flowing within reconstructed embryonic chick hearts. This pipeline avoids common bottlenecks by using clearing and imaging techniques that are cheap, fast, and easily customizable. In conclusion, this study demonstrates the utility of pairing advanced imaging techniques with computational models and GMMs by using light sheet fluorescence microscopy imaging to generate subject-specific CFD analysis in embryonic chick studies. The pipeline we’ve established can easily be modified and extended to explore various aspects of cardiovascular development in embryonic chicks.

## METHODS

### Chick Embryo Preparation

Fertilized White Leghorn (*Gallus domesticus*) eggs were incubated in a humidified incubator at 38°C to the desired developmental stages of approximately Hamburger-Hamilton (HH) 19^40^, which was approximately 2.5 days^41^. Development stages were confirmed using morphological features^42^. Each eggshell was windowed, and the shell membrane was removed to expose the embryo. 1-2 ml of phosphate-buffered saline was added to each egg and parafilm was placed over the window to prevent the embryos from dehydrating. Following these micro-injections, the embryos were removed from the eggs and placed in 6-well plates, and washed in Paraformaldehyde to arrest hearts. The embryos were shaken in methanol overnight at 4°C prior to clearing. Embryos were cleared using the iDISCO+ protocol with the alternative pretreatment and stored in glass vials prior to imaging.

### Imaging and Post-processing

Chick embryos were imaged at the UNC Microscopy Services Laboratory using the LaVision BioTec Ultramicroscope II^47^. The Imaris (Bitplane, Oxford Instruments)^48^ File Converter software was used to convert light sheet fluorescence microscopy-generated TIFF files to Imaris files.

### Computational Model

Grid spacing was approximately 0.009 mm with each embryo containing at least 100,000 second-order tetrahedral elements. Blood flow was assumed to be laminar^57^. Four cardiac cycles were completed for each simulation with a time step size of 0.01 s.

### Post Processing and Analysis

Simulations and resulting hemodynamics were visualized in Paraview (Kitware)^59^. Aortic arch centerline extraction was performed using the vascular modeling toolkit module VMTKSlicerModule (https://github.com/vmtk/SlicerExtension-VMTK) in the open-source software 3D Slicer (https://www.slicer.org/) to determine the length, tortuosity, curvature, average radii, and other geometric properties of each aortic arch. Radii were determined by calculating the maximally inscribed sphere along points in the arches. Geometric visualization and analyses were performed using RStudio (https://posit.co/)^61^ in R (https://www.r-project.org/)^62^.

